# Altruism pays off in group-structured populations through probable reciprocity

**DOI:** 10.1101/2024.01.20.575560

**Authors:** Klaus F. Steiner

## Abstract

The evolution of altruistic traits is a Darwinian puzzle that has not yet been satisfactorily solved. Using multi-agent simulations for public goods games and a threshold model, I show here that there is a clear fitness advantage for the altruists in a mixed population of altruists and egoists if the following conditions are met:

(i) The population is segregated into groups that persist over time or several generations. (ii) The benefit achieved by altruists is only shared among the members of their own group. (iii) There is selection pressure on each individual, which is reduced by the benefit provided by altruists. (iv) The costs for the altruists are smaller than the overall benefit they achieve for all members of their group together. (v) There is already a high proportion of altruists in the population and/or the proportion of altruists varies greatly between groups.

I refer this kind of population dynamics as the grouping effect, or G-effect for short. This effect is made up of two biases, which basically come about through simple individual selection: the tendency to increase the variance among the groups (*variation bias*) and the tendency to increase fitness for altruistic traits when this variance is high (*fitness bias*).

In this context, selection only happens at the individual level. It does not require relationships between the group members, nor does it require competition among the groups. Extended multi-agent simulations with reproduction, mutation and migration also show that altruism can exist stably in a wide range of conditions due to this G-effect and the resulting probable reciprocity.

## Introduction

The evolution of cooperation play a dominant role in biology, from the formation of simple cell colonies to human civilizations and the eusociality of ants and others. The general problem is that cooperation often means that you put needs aside on the one hand and run the risk of being exploited on the other.

In evolutionary biology, a trait that promotes cooperation is said to be altruistic if it benefits other individuals but costs its bearer more than it benefits them. Such a trait, which ultimately has a negative effect on the number of offspring and thus reduces the fitness of the carrier, is a so-called “Darwinian puzzle”. It represents a serious problem for the theory of evolution, which Darwin himself drew attention to (Darwin, 1888; Dugatkin, 2017; Okasha, 2020).

Several mechanisms have been proposed for solving the altruism puzzle and the evolution of cooperation, but there is no consensus on their validity. Nowak (Nowak, 2012) distinguishes between the following five mechanisms.

The most widespread approach in biology is kin selection, which relies on the genetic relationship of the cooperating actors (Hamilton, 1964). The main beneficiary of the cooperation is not the individual, but the common “gene” that the relatives share. An individual can therefore even sacrifice its life if this saves enough relatives to make it worthwhile for the common gene. Only the “inclusive fitness” must show a positive balance.

A different and often seen opposite concept is group selection. The basic idea goes back to Darwin’s assumption that groups compete with each other (Darwin, 1888). This means that groups of individuals, rather than individuals, are the units on which selection acts. A modern version of this, multilevel selection, assumes that selection acts at different levels, such as both the individual and the group level (Sober and Wilson, 1998; Wilson and Wilson, 2008).

Both the proponents of kin selection and those of multilevel selection often use the Price equation as their mathematical basis, as it indicates a predisposition to altruism (Frank, 1997, 1995; Price, 1970).

Two further solution mechanisms for the altruism puzzle are direct and indirect reciprocity. In the case of direct reciprocity (Trivers, 1971), the costs and benefits of repeated interactions between the actors are extrapolated and analyzed using game theory. In the case of indirect reciprocity, the alleged altruism is rewarded with reputation, which later benefits the actor. Both approaches attempt to eliminate the negative fitness balance of apparent altruism and show that it is in fact not an altruistic trait after all, but serves one’s own advantage.

The fifth mechanism for the evolution of cooperation is spatial selection (Nowak, 2012). This is based on the fact that not every individual can interact with every other individual, but that the population is spatially structured and interactions only take place in a limited environment.

In this paper, I show that the group-structured population may indeed be the key to the general solution of the altruism puzzle. On the basis of multi-generational public goods game models and cost-benefit calculations as well as multi-agent simulations, I demonstrate that altruistic traits in a group-structured population prevail purely arithmetically, given appropriate selection pressure on individuals, and that neither competition between groups nor kin preference is necessary for this. Like previous work on structured populations (Fletcher and Doebeli, 2009; Fletcher and Zwick, 2004; Killingback et al., 2006), this study emphasizes and proves the fundamental importance of spatial selection for the evolution of altruism.

## Models and results

In my models, I examine the costs and benefits of cooperative and altruistic traits in populations that are divided into groups. By groups, I mean parts of the population whose members share the benefits that cooperative members contribute to group welfare. This definition includes social groups and packs, as well as anonymous herds or island populations.

The populations consist of two types of individuals: cooperative or even altruistic cooperators C, who contribute to the welfare of the group at their own expense, and the “egoistic” defectors D, who contribute nothing to the welfare of the group and therefore have no costs.

The contribution of a C to the group is called benefit. It is important to distinguish between the individual payoff and the benefit for the group. In my models, the benefit is divided equally among all group members. Each member thus receives the same payoff: payoff = benefit / group size.

To keep the evaluations clear, I use ¢ (“cents”) as the unit for all costs, benefits and payoffs, and I use the term “scores” to refer to the individual ¢ values achieved.

A trivial win-win cooperation exists if the payoff of a C exceeds its costs, for example if C pays an amount of 2 ¢ and receives 3 ¢ as a payoff. This would be the case if C’s contribution to its group, consisting of itself and two Ds, generates a benefit of 9 ¢. The benefit for C is obvious here.

Altruism only exists if the costs for C are greater than its payoff, for example if C pays 2 ¢ and only receives 1 ¢ because the joint benefit for the same group of three is only 3 ¢.

Here I only examine evolutionary models without competition between individuals and groups. For this purpose, I define an objective, global survival threshold as the minimum score below which the individual is not viable. All above-threshold individuals, whether C or D, survive and reproduce with the same probability in the extended models.

The change in fitness of a type is equivalent to the change in its relative frequency. For the relative frequency of C in the population, I write [C], analogous to the notation for concentrations in chemistry. [C] is therefore the proportion of C in a population of C and D, and this is also the fitness of C in this population. [C’] stands for the frequency (and fitness) of C in a later state of the population. The change in fitness of the cooperative trait is therefore [C’] – [C].

The calculations are primarily based on multi-agent simulation models that I created with Netlogo (Wilensky, 1999).

### Selection models without reproduction

First, I examine pure selection events without reproduction, migration and mutation. So it is only about the elimination of individuals based on the respective scores achieved, i. e. whether the score reaches the threshold for survival or not. Unless otherwise stated, the following models use the values: costs *K* = 5 ¢, benefit *B* = 10 ¢ and threshold *T* = 3 ¢.

#### The simplest case: populations of two pairs

First we consider the possibilities for populations that consist of only two pairs. A pair can consist of two cooperators (CC), two defectors (DD) or one of each (CD). As the order does not matter, there are only 6 different combinations for populations consisting of two pairs: CC:CC, CC:CD, CC:DD, CD:CD, DD:CD and DD:DD. What happens to these six starting populations if the individuals with sub-threshold scores are eliminated? Three examples should illustrate the development (see also Fig. 1A).

**Fig. 1:**
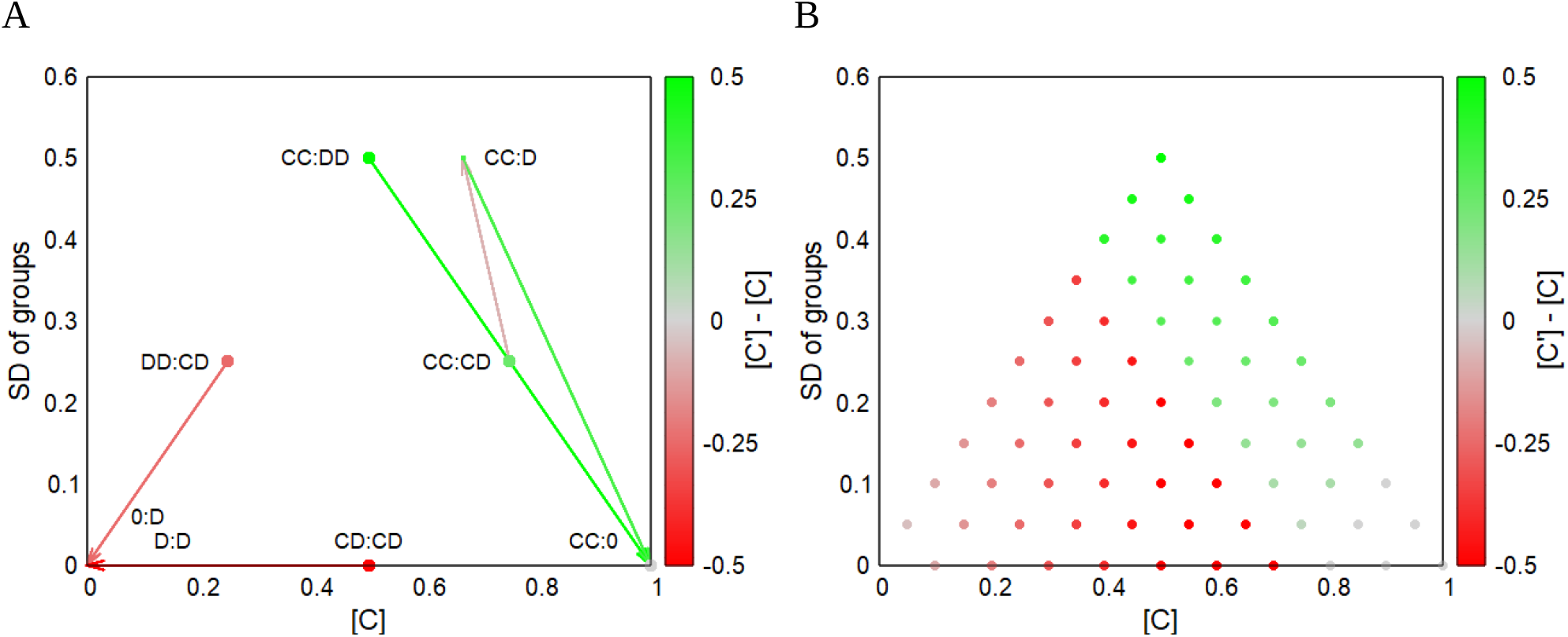
Fitness maps for two groups with 2 and 10 members each. Each dot represents a starting population, its color indicates the change in the respective proportion of C compared to the final population. The “greener” the greater the fitness gain, i. e. the higher the proportion of C in the final population. A: Fitness map with trajectories for starting populations of 2 groups with 2 members each (see also examples described in the text). The color of the trajectory corresponds to the fitness change between consecutive steps. B: Fitness map for the 66 possible starting populations of two groups with 10 members each. The colors of the dots again indicate the fitness differences to the final population. Here 30 of the 66 starting populations lead to final populations with a high C-ratio [C], the others lead to the extinction of the population. The trajectories are not shown here.

##### Example A

The starting population **CC:DD** consists of a pure C and a pure D group with two individuals each. The frequency of the C in the population is [C]=0.5 and the standard deviation between the groups is SD=0.5 (SD is the standard deviation of the frequencies [C_group1_]=1 and [C_group2_]=0). The benefit from the two Cs in the first group is 20 ¢, that is 10 ¢ for each. Because they are Cs, a cost of 5 ¢ is deducted from each of them, making a score of 5 ¢ for each C in the first group. In the second group of 2 D, there is no benefit and no cost, making a score of 0 ¢ for each. Since the threshold is 3 ¢, the Ds are eliminated and both Cs survive. The new group composition is **CC:0**, so one group of 2 Cs remains and the second is now “empty”. There are only Cs left, therefore [C]=1 and the SD=0. A pure C population remains.

##### Example B

In the initial population **DD:CD**, the first group consists of 2D and the second of 1C and 1D, the frequency of C is thus [C]=0.25 and the standard deviation of groups is also SD=0.25 (SD of [C_group1_]=0 and [C_group2_]=0.5). The two Ds in the first group again have a sub-threshold score of 0 ¢ and thus die. The second group has a benefit of 10 ¢ from the one C, which is divided among the members. D thus achieves a score of 5 ¢, is therefore above-threshold and survives. The costs of 5 ¢ are deducted from C and it therefore has a sub-threshold, lethal score. The new population is therefore 0:D, there is now only one group, one D and no C, that is [C]=0 and SD=0. The lone D then dies in the next round with 0 ¢, which means that the population is extinct.

##### Example C

In the starting population **CC:CD** we have [C]=0.75 and SD=0.25 (SD of [C_group1_]=1 and [C_group2_]=0.5). The Cs in the first group thus score 5 ¢ each and survive, in the second group D also scores 5 ¢ but C gets nothing and dies by falling below the threshold. The new population is thus **CC:D** ([C]=0.67 and SD=0.5). The two Cs in the first group thus achieve 5 ¢ again. The D is now alone in the second group, is therefore left empty-handed and does not make it any further.

Ultimately, the population is **CC:0** ([C]=1 and SD=0, as there is only one group left). This leaves a pure C population again.

For the sake of completeness, here are the other possible starting populations: **CD:CD** ([C]=0.5 and SD=0) leads to **D:D** and in the next step, like **DD:DD** ([C]=0 and SD=0), to complete extinction. A pure C population **CC:CC**, on the other hand, remains unchanged at [C]=1 and SD=0.

#### What would have happened to the starting populations if they had not been divided into groups?

Spoiler: All three start populations from the examples would perish and not just the one from example B.

The initial population **CC:DD** from example A are 2C and 2D. Without grouping, the four would share the 20 ¢ benefit from the two Cs, making 5 ¢ for each D and only 0 ¢ for the Cs because of their costs. Cs are sub-threshold and die. This leaves only the 2 Ds, which are eliminated in the next round with 0 ¢ each.

Start population **DD:CD** from example B consists of 1C and 3D. The four share the benefit of 10 ¢ from C, making 2.5 ¢ for D and -2.5 ¢ for C. Everyone is sub-threshold and the population dies out.

In example C with **CC:CD**, the benefit is 30 ¢, making 7.5 ¢ for D and, after deducting the costs, 2.5 ¢ for each C. The Cs are therefore sub-threshold and disappear. D survives, but perishes in the next round with 0 ¢.

An unstructured population of four could only exist under these conditions if all four are C (**CCCC**), only then are they all above threshold with 5 ¢ each.

#### Larger and more groups

If larger or more groups are involved, the number of possible starting populations increases according to an urn problem with replacement and where order does not matter. Fig. 1B, for example, shows the fitness map for all 66 possible starting populations consisting of two groups of 10 individuals each. The principle remains the same. From the starting population, a few intermediate steps lead to a fixed final state either with a high [C] or without C, which is equivalent to the extinction of the population. Fig. 2 shows a concrete example of the development of a starting population (A) consisting of 16 groups of 10 individuals each, which freezes in the final state (C) via an intermediate step (B). The relative fitness of the cooperators [C] increases from the original 0.719 to 0.833, meaning that the altruists become fitter compared to the defectors. In absolute terms, their number decreases from 115 to 75 and “empty groups” also arise, but from an evolutionary point of view only the ratio of potential replicators is relevant.

**Fig. 2:**
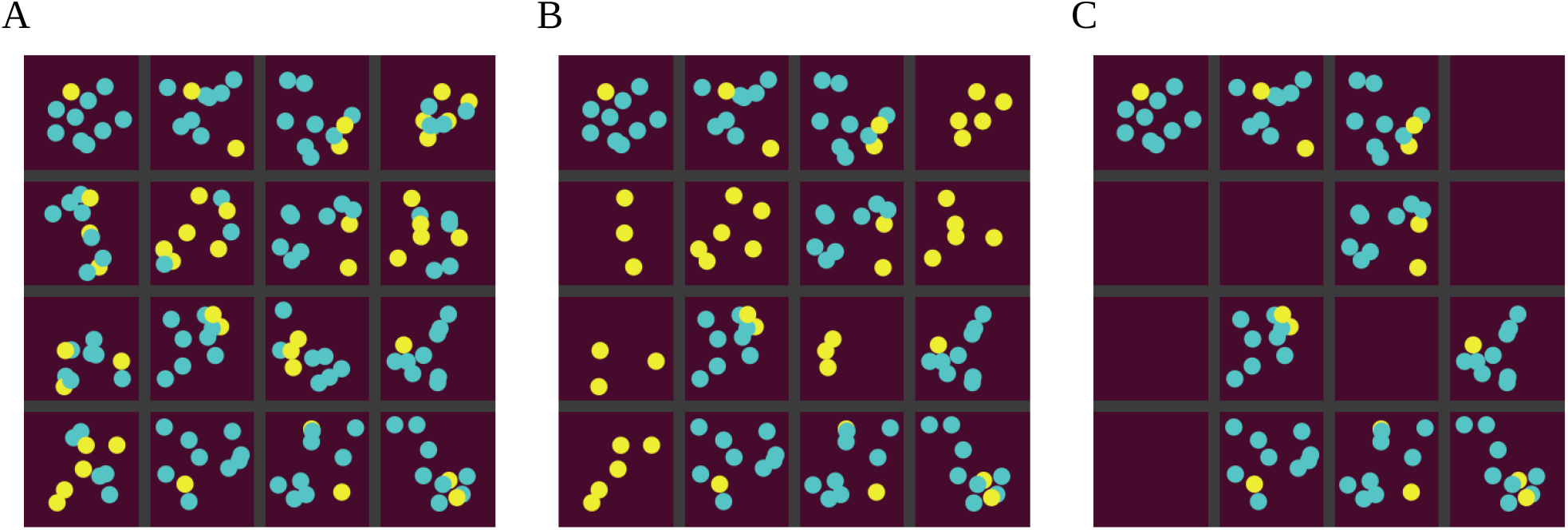
Development of a starting population of 16 groups with 10 individuals each. Blue dots are cooperators (C), yellow dots are defectors (D), the gray lines separate the groups. A: Random starting population with a total of 115 C and 45 D ([C]=0.719, SD=0.155). B: Situation after the first selection step (75 C, 45 D, [C]=0.625, SD=0.415). C: Final state after the second selection step (75 C, 15 D, [C]=0.833, SD=0.047). All individuals are now above the threshold, so nothing changes and the final population is “frozen”. The relative fitness of C has thus increased by 0.114 between the start and end population.

Fig. 2A shows one of the 5,311,735 possible starting populations consisting of 16 groups and 10 members each. The fitness map in Fig. 3B shows the fitness changes of 1000 of these starting populations and their final populations. Five of these are shown in Fig. 3A together with their trajectories to their respective final populations. Here, in the two starting populations with the lowest [C], the C die out immediately, so [C’]=0 and there is also no variance (SD=0). The fitness of the C therefore decreases ([C’]-[C]<0), accordingly the starting populations are colored reddish. (Since D alone is always sub-threshold, the entire population perishes. However, this is not directly visible in the fitness map).

**Fig. 3:**
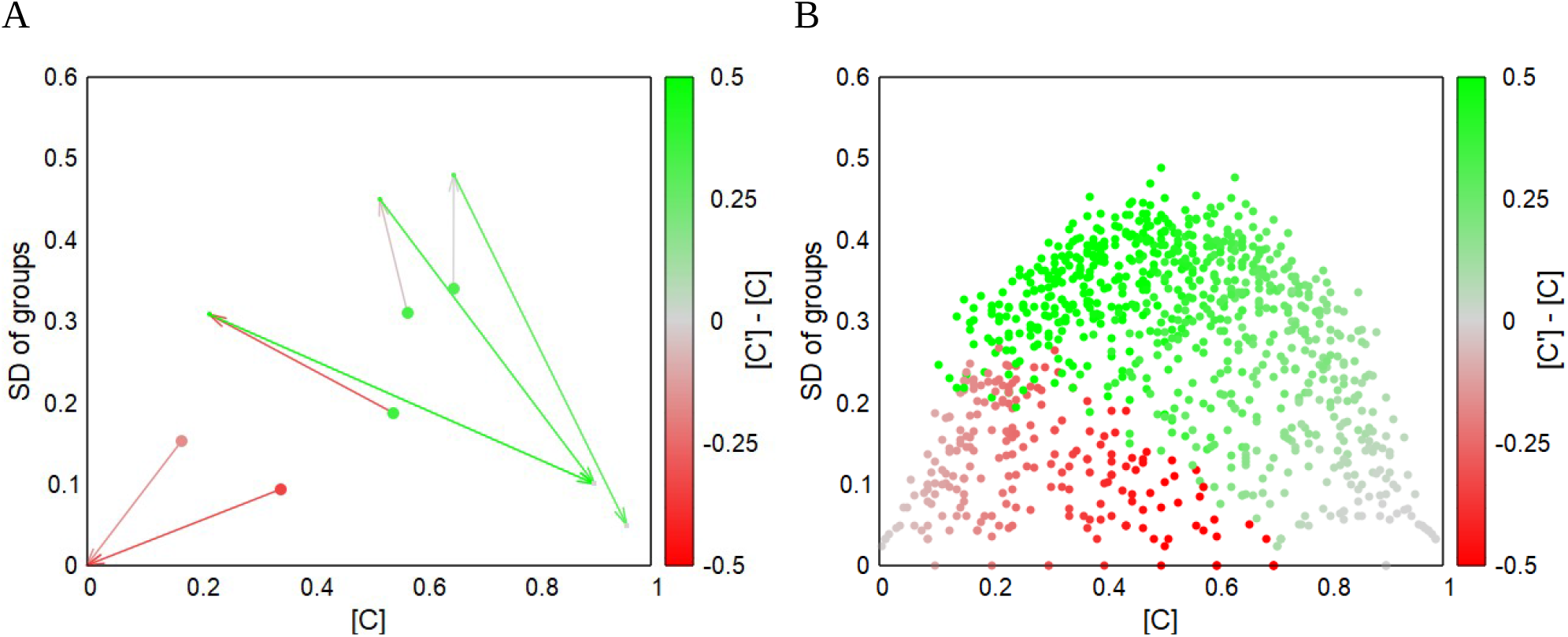
Development of populations consisting of 16 groups of 10 individuals each. A: Trajectories of five random starting populations. B: Fitness map for 1000 starting populations and the changes in [C] (=fitness) compared to their final populations. 77.8 % of these starting populations end up in stable populations with a high proportion of C (green), the remaining 22.2 % die out (red).

The three greenish starting populations in Fig. 3A achieve a gain in fitness up to the final population, i. e. a relative increase in C. In one case there is initially a marked decrease in fitness, but in the final state all three are positive. A second component is noticeable in all three cases: the sometimes significant increase in variance after the first iteration.

The change in fitness is only half the battle. Just as important is the change in variance (SD of groups) in order to be able to describe the development of a population. Fig. 4A shows the fitness map for the first iteration and Fig. 4B the corresponding “variation map”, in which the expansion of the trajectories on the y-axis is color-coded. For blue starting populations the variance increases, for red it decreases and for gray there is little or no change. Note that the increase in variance (blue dots in 4B) is more pronounced than the increase in fitness (green dots in 4A).

**Fig. 4:**
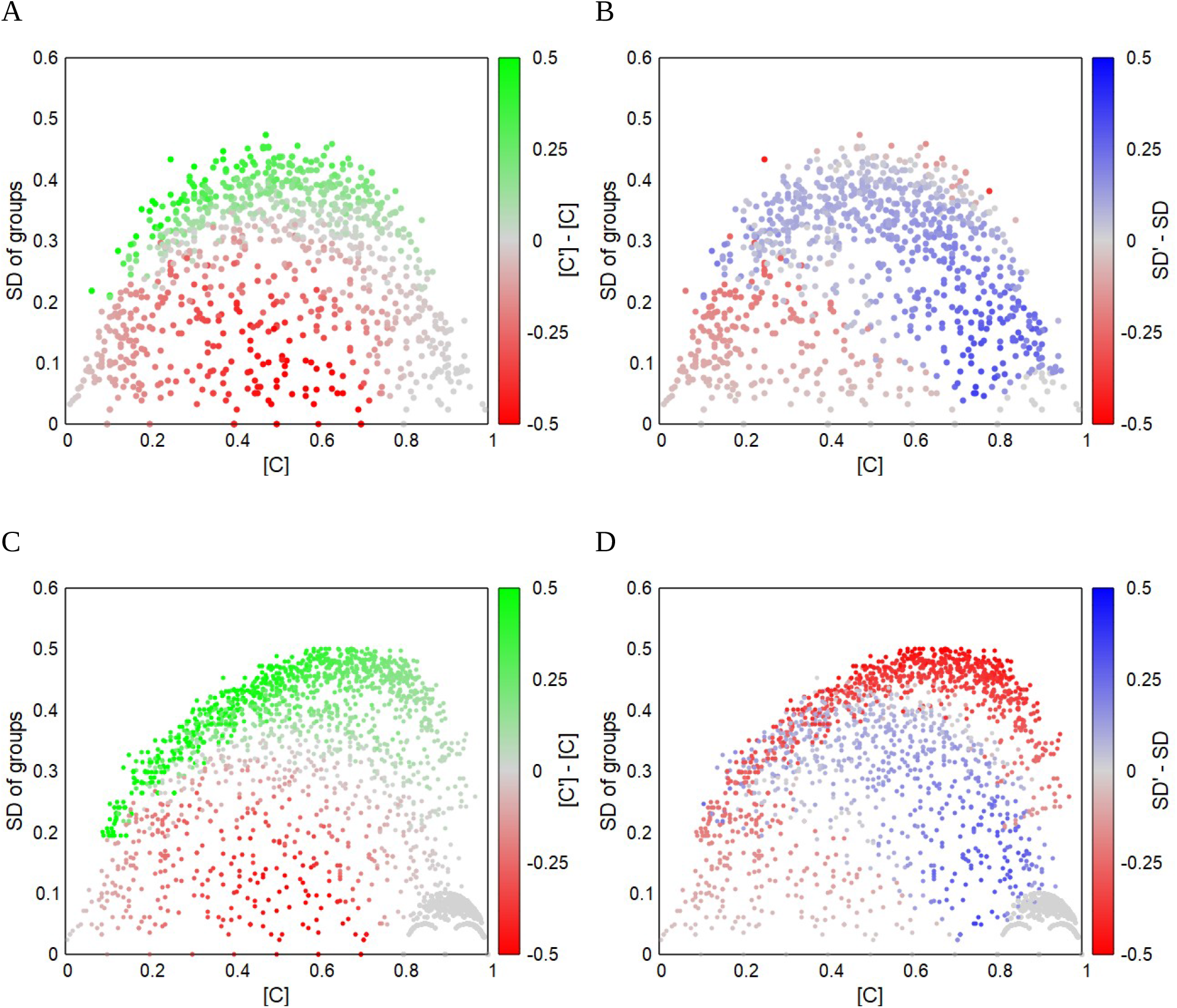
Fitness and variation maps for the development of 1000 starting populations consisting of 16 groups of 10 individuals each (*K*=5, *B*=10 and *T*=3). A: Change in fitness after the first selection step. B: Change in the standard deviation of the groups after the first selection step. C: Change in fitness of all start, intermediate and final populations. D: Change in the standard deviation of the groups of all start, intermediate and end populations (variation map). In the gray areas at [C]>0.8 and SD<0.12 the stable, “frozen” final populations are collected.

Fig. 4C and 4D also show the fitness and SD changes of all intermediate populations. In contrast to the starting populations with always 16 groups of 10 members, individuals or even entire groups have usually been lost from the subsequent populations, which means that completely different [C] to SD distributions are possible.

Many of the starting populations in 4B show a trend towards very high variances. These intermediate populations with very high SD in turn show a clear tendency to increase fitness (green in 4C) and decrease SD (blue in 4D) and thus end up in the “frozen zone” with the stable final populations.

#### The threshold to altruism

So far, we have only considered threshold models with costs *K* = 5 ¢, benefits *B* = 10 ¢ and threshold *T* = 3 ¢ and we have seen that altruism can persist as soon as the population is divided into groups. Let us now look at the development of populations with other costs, benefit values and thresholds. The interesting case here is when the benefit for the group is greater than the cost for C, that is *B > K*. The influence of the threshold value under these conditions is shown in Table 1 and the fitness maps in Fig. 5B-F.

**Table 1:** Development of the population at different threshold values *T*, if B > K.

| Threshold $T$ | Effect | Dominance |
| --- | --- | --- |
| $T \leq -K$ | No selection pressure on C and D, therefore starting population already freezes (Fig. 5B) | $C \approx D$ |
| $-K < T \leq 0 \text{ ¢}$ | Some selection pressure only on C, as only C can have scores $< 0 \text{ ¢}$. Below-threshold C are eliminated until groups become fixed (Fig. 5C) | $D > C$ |
| $0 \text{ ¢} < T \leq B - K$ | Selection pressure on C and D, but C are relatively fitter than D, as shown in this paper. Range of the G-effect and stable altruism (Fig. 5D and E as well as Fig. 1-4) | **C** |
| $B - K < T$ | Lethal selection pressure on C and D. Extinction of the population (Fig. 5F) | † |

**Fig. 5:**
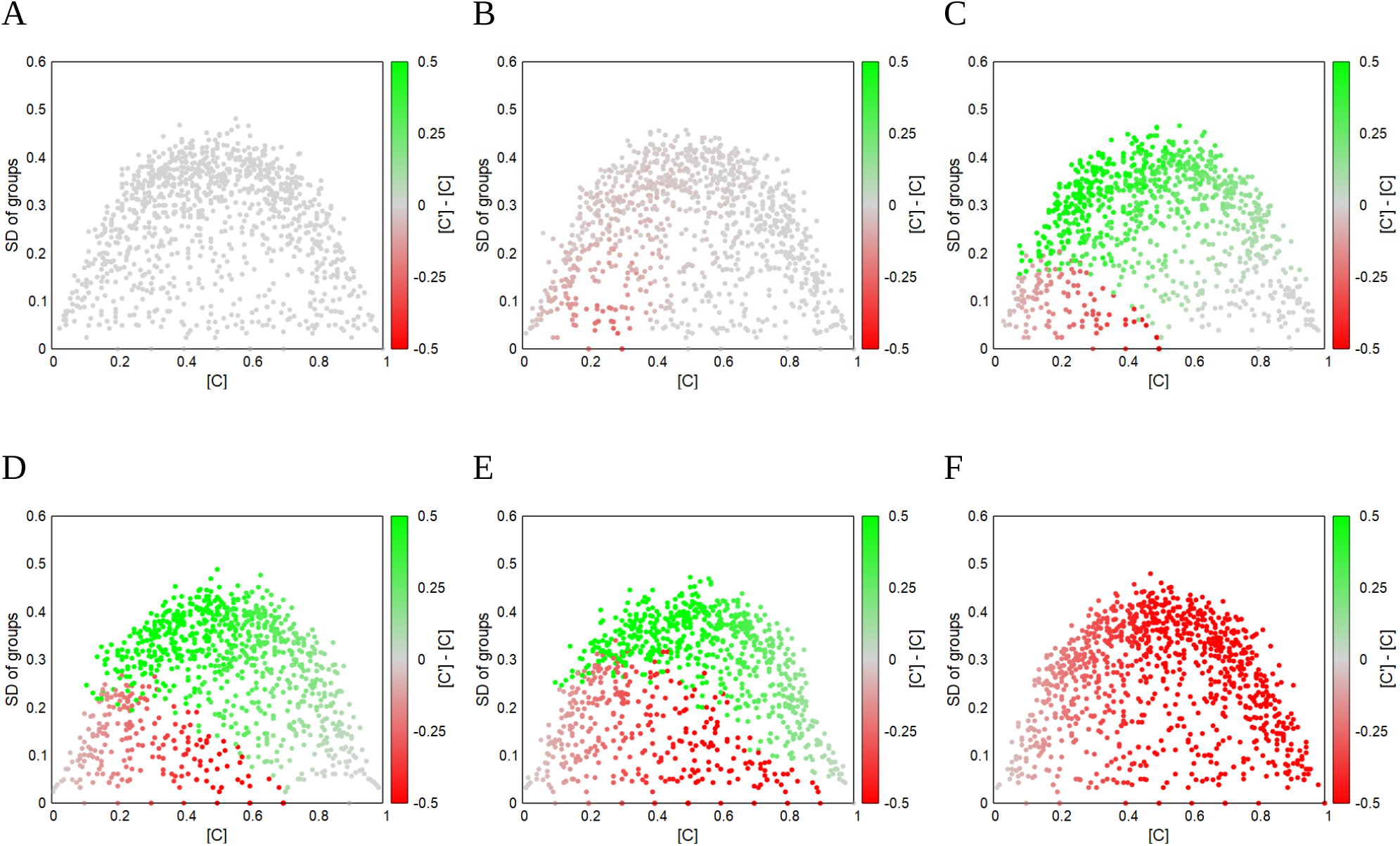
Fitness maps for different thresholds when *K*=5 and *B*=10. A: Threshold *T* = -6 means that D and C are always above threshold and therefore there is no selection pressure for either. B: At threshold *T* =-1, only C can be sub-threshold at low [C], therefore there is slight selection pressure for C and none for D. C: Threshold *T* = 1 favors altruism and fitness gain for C over wide [C] ranges. 87.5 % of all start populations end up as populations of only or mainly C. D: Same map as shown in Fig. 2B with *T* = 3, where 77.8 % ended up as C-populations. E: At threshold *T* = 5, stable altruism is still possible, but there are more starting populations that lead to extinction than at *T* = 1 or *T* = 3, so only 64.5 % remain as C-populations. F: At threshold *T* = 6, C and D are always sub-threshold, the entire population dies out immediately. All maps show the development of 1000 starting populations consisting of 16 groups of 10 individuals.

Stable altruism is therefore only possible if the threshold lies between 0 ¢, that is a pure D population ([C]=0), and *B – K*, which corresponds to a pure C population ([C]=1).

However, if the benefit for the group is less than (or equal to) the cost for C, i. e. if *B* ≤ *K*, stable altruism cannot establish itself. If the threshold *T* is in the range *B – K < T* ≤ 0 ¢, there is selection pressure only on C and therefore D prevails. If the threshold is higher (*T* > 0 ¢), C and D are sub-threshold and both die out.

### Models with reproduction, migration, mutation and additional mortality

So far, we have only considered selection alone, without reproduction, migration or additional mortality. If they do not die out, the populations therefore freeze in their final states after a few iteration steps and nothing more happens.

Adding reproduction, migration between groups and additional mortality factors to the model results in more dynamic systems. This paper cannot go into the many possible influencing factors, so only a brief overview of how such “more real” populations develop could be given here. For this purpose, I define a “standard model” in which I make changes to individual parameters and compare the effects on the stability of the populations, or more specifically: how many of 100 random starting populations still exist after 1000 generations.

As a **standard model**, I use a starting population of 16 groups of 10 individuals each, whether of type C or D is again chosen at random. In each group, the individual scores are calculated as before, e. with the parameters benefit *B* = 10 ¢ and cost *K* = 5 ¢, the threshold is again *T* = 3 ¢. Scorers below the threshold are eliminated at 100 % (*sub-threshold mortality*). In addition, 10 % of randomly selected individuals are eliminated regardless of their scores (*general mortality*). Each surviving individual produces two offspring for its group that deviate from the type of the parent with a probability of p=0.02 (*mutation rate*). Finally, in each generation, 10% of the individuals migrate to another group at random (*migration rate*).

The procedure for each starting population:

~~~
create random starting population of **number-of-groups** groups with **group-members**
individuals each repeat 1000 times:
calculate score for each individual based on benefit *B* = 10 and cost *K* = 5
eliminate with probability **sub-threshold mortality** all individuals with scores below the threshold *T* = 3
eliminate random individuals according to the **general mortality**
generate two offspring from each survivor for the same group, taking into account the **mutation rate**
eliminate surplus group members at random
let random individuals change to random other groups according to the **migration rate**
eliminate surplus group members at random again
If no more individuals exist, increase the counter for the extinct population, otherwise for the surviving population
~~~

#### Interpretation of the results

Like the selection models, the developments of these more complex models can also be depicted in fitness maps. Fig. 6B shows the trajectories of two random starting populations of the standard model. The red dot represents a starting population in which all C are immediately eliminated ([C]=0). Since pure D groups are not viable, the entire population dies out. As a reminder, without C in the group, each D has a score of 0 ¢, but the threshold is 3 ¢.

**Fig. 6:**
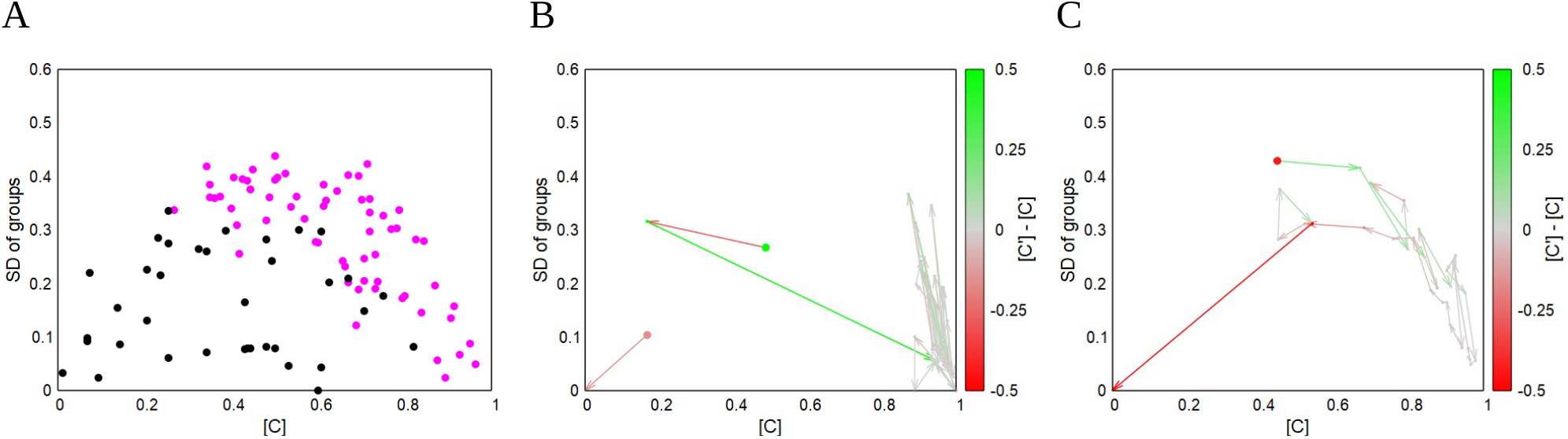
Survival and extinction of populations. A: Survivability of 100 starting populations in the standard model. Black dots show extinct starting populations, pink dots are starting populations that still exist after 1000 generations. Specifically, 64% of the starting populations survive. B: Example of two starting populations in the standard model, one of which (red dot) dies out immediately and the other (green dot) is stable and still exists after 1000 generations. C: Trajectory of a population with 30% migration that dies out after 31 generations.

The green starting population in Fig. 6B shows a long trajectory that remains in a high [C] range over 1000 generations. This population is and remains stable with over 90% altruists. Fig. 6B shows the survival or extinction of 100 starting populations from the standard model. If a population survives 1000 generations, the corresponding starting population is shown as a purple dot, and as a black dot if it dies out before then. Here, in the standard model, 64% of the starting populations still exist after 1000 generations. This result for the standard model is displayed as reference in all diagrams in Fig. 7 as the red bar. The blue bars show the survival rates of populations that deviate from the standard in exactly one parameter.

**Fig. 7:**
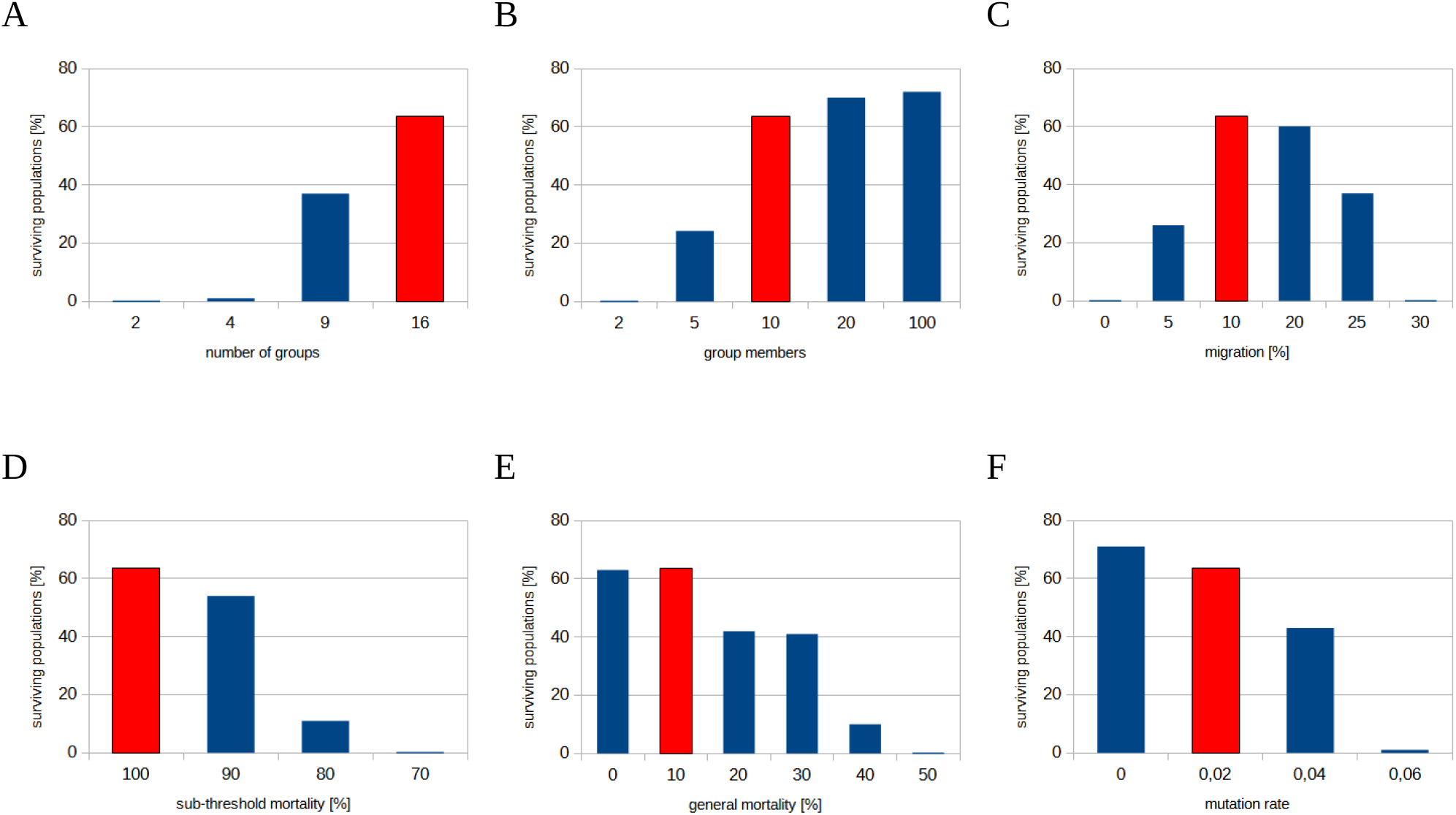
Frequency of random start populations that still exist after 1000 generations. The red columns show the results of the standard model, the blue columns show the results of models that deviate from the standard in exactly this one parameter. A: Effect of the number of groups (with 10 members each) for the stability of the population. B: Influence of group size with 16 groups. C: Migration among groups. Proportion of individuals that randomly move to a different group in each generation. D: Sub-threshold mortality is the probability that an individual with a score below threshold *T* will die. E: General mortality is the probability of death independent of the score. F: Mutation rate is the probability that a C has a D as offspring and vice versa. See text for details.

A brief interpretation of the results shown in Fig. 7 when varying the individual parameters:

#### Number of groups

The fewer the groups, the greater the probability that the population will die out. If there are only a few groups, it is more likely that all groups will suffer at the same time, this means that there will be relatively few Cs in each group and little variance between the groups. This leads to lower G-effects and the population dies out.

#### Group members

Populations composed of very small groups are more likely to die out than populations with larger groups. A smaller population size leads more frequently to “groups” with only one member, especially through migration, as it is less likely that at least two will immigrate into an empty group at the same time. However, “individual members” are always sub-threshold and are eliminated.

#### Migration

Migration is not directed in the models, there is also no preferential filling of empty or small groups and occurs after reduction to the maximum group size. Without migration, extinction occurs, as all groups eventually perish by chance due to general mortality and mutability.

Low migration makes the formation of new groups more difficult, as at least 2 members are required, because both C and D are sub-threshold on their own. In models where the surplus of offspring specifically fills other groups, even small populations divided into a few groups are very stable. However, this is material for future work.

Too much migration between groups, on the other hand, reduces the variance between groups. This reduces the G-effect and thus increases the extinction of the entire population.

#### Sub-threshold mortality

So far, we have always assumed 100% mortality of individuals with sub-threshold scores. However, the G-effect is also effective with selection pressures that are not absolutely lethal, albeit to a decreasing degree. In the standard model, for example, it is already too low at 70% to allow populations to exist over 1000 generations.

#### General mortality

In real situations, of course, it is not only the absence of altruists that causes mortality. An additional mortality of the same magnitude for C and D also prevents the “freezing” of groups in the model if all members are above threshold and thus ensures more dynamics. A very high general mortality leads to smaller populations, more empty groups and a higher probability of extinction.

#### Mutation rate

The mutation rate indicates the probability of an offspring being of an other type as its parent. With increasing mutation rate, the variance between the groups decreases on the one hand, but it also lowers high [C], since with high [C] the C also have more D as offspring. Both lower the G-effect and increase the probability of the population going extinct.

The results of these more complex models show the limits and at the same time illustrate the wide range over which stable populations can exist that are based solely on the benefits of altruistic individuals for their groups.

## Discussion

The simulation models presented in this paper demonstrate that altruism can exist by itself when populations are divided into multi-generational groups and selection pressures are just right; that under a wide range of conditions, stable populations can develop in which many altruists and a few egoists are in a dynamic equilibrium, where individual groups disappear and new ones emerge over the generations, but the populations still exist after a thousand generations.

I refer to the underlying processes as the grouping effect, or G-effect for short. It is made up of two steps that come about solely through individual selection.

*Step 1: Increasing the variance between the groups (variation bias)*

If, by chance or for other reasons, there are so many egoists (D) in a group that the altruists (C) become sub-threshold due to their extra costs, this quickly leads to an (almost) pure egoist group. If the other groups consist mainly of altruists, the variance between the groups is suddenly increased. This “variation bias” is shown in Fig. 4B and D.

*Step 2: Relative increase in fitness for altruists when variance is high (fitness bias)*

If the group now consists primarily of egoists, they are now also sub-threshold and eliminated in the next step. Their disappearance increases the relative frequency of altruists in the population globally. This “fitness bias” is illustrated in Fig. 4A and C.

The Price Equation appears to express only the fitness bias and does not take into account the effects of the variation bias. This is the reason why the tendency towards altruism described in this work is far stronger than the predisposition calculated by the Price equation alone (Frank, 1995; Grafen, 2000; Van Veelen, 2020). However, this topic requires further investigation.

### Conditions favoring the G-effect

In my model, there are some prerequisites for stable populations due to the G-effect:

1. The population is divided into permanent, multi-generational groups, with only limited migration between them. The population must be divided into groups that exist over a long period of time or several generations. If groups are soon mixed again, as in Wilson’s trait group model (Wilson, 1975), any variation that may have developed is lost and the “variation bias” does not come into play.
2. Benefits are shared among all group members. The benefit achieved by altruists is shared exclusively among the members of their own group. In my model, the benefits of several altruists are added up and distributed equally among all group members. As the donor also receives his share, we are dealing here with whole-group altruistic traits (Pepper, 2000).
3. Global selection pressure on individuals, which is mitigated by benefits. The selection pressure on each individual is reduced by the benefits contributed by altruists. The selection pressure on individuals comes from outside the group and is the same for all individuals in the population. It is best to think of harsh environmental factors that the individual cannot survive without others.
4. The costs for the individual altruist are lower than the benefit of his contribution to the group. The costs for the altruists are lower than the total benefit they achieve for all members of their group. Nevertheless, this is (mostly) altruism, as the benefit provided in is shared among all members of the group and the share for the altruists themselves is therefore generally smaller than their costs. I will discuss this in more detail below.
5. There are more altruists than defectors in the population or the variance between the groups is high. There is already a high proportion of altruists in the population and/or the proportion of altruists varies greatly between the groups. Too few altruists and little variance means that the proportion of altruists is also too low in all groups. As a result, the altruists in all groups become sub-threshold at the same time, followed by the egoists in the next step, and the entire population becomes extinct.

### Three perspectives

This simple multi-agent model with selection thresholds already shows elementary sociological phenomena, which I would like to look at briefly from different angles: from the perspective of the individual, the group and the population. I will use the dynamic properties of my “standard model” for the discussion.

#### From the perspective of the individuals: altruists and egoists

An altruist C in a group with a high proportion of altruists receives sufficient benefit from the others so that its costs are not significant and there is no or only slightly increased selection pressure for this individual. An egoistic D in the same group may have advantages on paper (higher scores), but they have no effect on its reproduction, as all group members are above-threshold and therefore everyone in the group is “doing well”.

However, once the selfish defectors D get out of hand in the group, C soon becomes a victim of its expensive altruistic trait. The resulting disappearance of C in the group soon causes the scores of D to drop to such an extent that they too become subliminal and die out. The “exploitation” of the C only affects the D with a delay in the coming generations, but it affects them just as fatally.

#### From the groups’ point of view: prosperity and decline

A group with a high proportion of C players is prospering. All members have scores above the threshold, single defectors D do not really harm the others. Under favorable circumstances, the probability of this group remaining in this state is high. Members migrating from these thriving groups are primarily altruists themselves.

However, if, by chance or due to unfavorable circumstances, the number of Ds reaches a critical value in a group, the group can quickly go downhill due to positive feedback. First the Cs die, which become subliminal because of their extra costs. As a result, only the D survive and reproduce, but without the benefits of the C, they soon fall below the threshold and also perish.

This decline may be cushioned by immigrating Cs from prospering groups. If this is not enough and the group dies out, a new group is soon formed by the influx of these mostly altruistic immigrants, who can then build up a prosperous group again.

This social cycle of prosperity and decline of groups is thus subject to an intrinsic dynamic.

#### From the populations’ perspective: homeostasis and extinction

If you look at a population over generations, you will mainly see altruists (C) and far fewer egoists (D) appearing and disappearing again. Time and again, however, individual groups show a sudden proliferation of egoists, often followed by the extinction of the entire egoist group.

Individual egoist groups suddenly increase the variance of the groups in the population, and the subsequent extinction of the egoist group in turn increases the relative frequency of altruists in the population, and so their fitness increases. Both together form the G-effect, which I have presented in this paper.

This creates a kind of homeostasis at the population level, which maintains a stable C:D or altruist to egoist ratio with strong advantages for C, as if controlled by an invisible hand.

If, on the other hand, too many groups decline synchronously, the G-effect is no longer sufficient and the whole population dies out. Over a wide range of parameters (see Fig. 7), however, the standard model shows a dynamic equilibrium of many prosperous and few declining groups.

## Conclusions

Finally, I will briefly discuss the core elements and the G-effect in the context of kin and group selection.

### Thresholds and the level of selection

A global threshold as selection criterion is best understood as an abiotic factor that is the same for all individuals in the population. Using a global threshold means that the “level of selection” in the model is explicitly the individual. Although the G-effect is based on groups, it does not require any selection pressure on these groups, as demanded by representatives of group selection.

A threshold model also has the advantage that it does not become entangled in actual or supposed statistical pitfalls, such as the Simpson’s Paradox or the Averaging Fallacy (Sober and Wilson, 1998), as selection is not based on averaged values. Instead, each individual’s score is measured against a fixed threshold value.

### Altruism is “relative”

The cooperators C are not always altruists, even if the costs, benefits and threshold remain the same. Whether it is altruism or not depends on the others.

The main reason for this is “wealth”. In a group consisting of almost only C and in which all members, regardless of whether C or D, are well above the threshold, there is no additional selection pressure for C due to the costs. The costs are therefore a purely marginal problem here and of no significance for the survival of the individuals.

A second, minor reason why C aren’t always altruistic in my models is that I have considered whole-group altruistic traits where the benefit is shared equally among all group members. If the group is very small and the benefit for the group is high, the payoff for C can be greater than its costs. Thus C makes a direct profit from its own trait, if the costs are less than the benefit for the group divided by the number of group members (*K<B/N*). With others-only traits, where C does not get his own share back, this effect would not occur (Fletcher and Zwick, 2004; Pepper, 2000).

The C individuals are therefore more likely to be “potential altruists”, although each individual has a completely fixed program. The same trait can be altruistic at one time and not at another, depending only on the number and type of group members among whom the benefit is shared.

#### Evolution of altruism

As described, starting populations with predominantly altruists are a prerequisite for stable “altruistic” populations due to the G-effect. Accordingly, a single altruist has no chance of gaining a foothold alone among egoists in my standard model. Nevertheless, I found two scenarios in which an altruistic trait can be established in a purely selfish starting population.

In the first approach, there is initially no selection pressure on C and D, i.e. the “altruistic” trait is neutral, or in the context of the model when *T* ≤ -*K (*or T ≤ 0 if *K<B/N)*. If there is only little migration between the groups, this automatically results in large variance between the groups, because almost pure C and D groups are created by chance and inheritance. This gives a favorable starting situation for altruism: high variance between groups. If the selection pressure now increases so that the threshold lies in the range 0 ¢ < *T* ≤ *B - K*, a stable population with predominantly altruists quickly develops due to the G-effect.

For the second approach, it is crucial that the sub-threshold mortality rate is less than 100% and here, too, restricted migration is important. It works best when empty groups are filled by migration and there is otherwise little exchange between existing groups. If the sub-threshold mortality is low, the altruistic trait cannot establish even if threshold level would allow this (0 ¢ *<* ***T*** ≤ *B – K*). Above a certain sub-threshold mortality rate, however, the altruistic trait suddenly establishes itself and then remains even if the sub-threshold mortality rate is increased to 100%.

Of crucial importance for the evolution of altruism in both scenarios is limited migration between the groups and an increasing selection pressure on all individuals (C and D), whether by shifting either the threshold or the sub-threshold mortality. This finding supports old assumptions that cooperation, mutual aid and altruism occur especially under harsh environmental conditions (Kropotkin, 1902). Both scenarios can occur with large groups, so it could also be relevant for the evolution of altruistic traits in bacteria and other microbes (Cremer et al., 2019).

I have only briefly outlined two promising scenarios for the evolution of altruistic traits here, further research is needed.

### What about kin and group selection?

This work shows that the G-effect is sufficient for the evolution of stable altruistic traits. It does not require kinship between group members, as greater relatedness within the groups is rather the result of the sorting caused by the G-effect and limited migration. Nor does it require competition between groups, as postulated for group selection.

The G-effect forms the basis for altruism and cooperation in any group structured population, and presumably in any large population, as it is structured by the different distances between individuals alone. This effect can be supported by competition between the groups as well as by favoring relatives, as both can have impacts on individual selection pressure. However, neither is necessary for the evolution of altruism, as previous work on group-structured populations has already pointed out (Fletcher and Doebeli, 2009; Fletcher and Zwick, 2004; Killingback et al., 2006).

The altruism in the population is promoted by the probable reciprocity that is supported by the structuring of the population and the resulting G-effect. Therefore, altruistic individuals “bet” with good odds that there will be mostly other altruists in their group. Some, however, are unlucky and find themselves in groups with too many exploitative defectors. These are the pawn sacrifices of their altruistic strategy. Thus, this work suggests that an evolutionary poker game solves the Darwinian puzzle of altruism.

## Used tools

For language support, I regularly used DeepL (https://www.deepl.com) and Google Translator (https://translate.google.com), because my native language is German. I also used Microsoft Clipchamp to create the demonstration video.

Otherwise, I didn’t use any AI tools, especially not for creating program code.

I used NetLogo, Python and LibreOffice Calc for the modeling and calculation tasks and Gnuplot for the visualizations. I wrote all the program code myself.

Used software:

- NetLogo, as programming environment for multi-agent-models: http://ccl.northwestern.edu/netlogo/
- Python, for scripting in-between: https://www.python.org
- LibreOffice, for writing (Write) and subsidiary calculations in Calc: https://de.libreoffice.org/
- Gnuplot, for the creation of the diagrams: http://gnuplot.info

## Supplemental material

An online version of my simulation program and a short video demonstrating the most important features can be found at https://www.vinckensteiner.com/ksteiner

